# Piloting floral-visitor assemblages: Camera-based monitoring in the cliff-dwelling Eastern Alpine endemic *Physoplexis comosa* (Campanulaceae)

**DOI:** 10.64898/2026.10.01.756015

**Authors:** Katerina Iberl, Philipp Kirschner, Alptekin Koç, Tereza Hasáková, Camilla Wellstein, Jakub Štenc

**Affiliations:** Department of Botany, Faculty of Science, Charles University, Prague, 128 41, Czech Republic; Institute of Botany, University of Innsbruck, Austria; Faculty of Agricultural, Environmental and Food Sciences, Free University Bozen-Bolzano, Italy; Department of Zoology, Faculty of Science, Charles University, Prague, 12800, Czech Republic; CREAF - Centre de Recerca Ecològica i Aplicacions Forestals, 08193 Bellaterra, Barcelona, Spain

**Keywords:** Biotic interactions, floral visitors, action cameras, Alpine endemics, *Physoplexis comosa*, eDNA metabarcoding comparison

## Abstract

Understanding floral visitor dynamics along elevational gradients is essential for assessing potential implications for pollen limitation and population genetic processes in fragmented populations of cliff-dwelling alpine endemics. *Physoplexis comosa* (Campanulaceae) is a chasmophytic paleoendemic restricted to vertical rock faces on dolomitic and limestone bedrock from 200 to 2,800 m a.s.l. The species exhibits a specific floral morphology with fused corolla apices, basal openings for nectar access, and secondary pollen presentation, yet its functional floral visitor assemblage has remained poorly characterized across the species’ elevational range.

In this pilot study, we tested a two-tier camera-monitoring setup combining action cameras for diurnal monitoring with Raspberry Pi-based systems equipped with IR illumination for combined diurnal and nocturnal recording. We conducted monitoring across five elevational sites (1,480–2,400 m a.s.l.) in the Eastern Alps. Recorded video material was downsampled and annotated using the Addax platform alongside manual cross-validation to record physical contact events.

Preliminary results confirm the technical feasibility of continuous monitoring in steep terrain and suggest possible elevational variation in diurnal visitors, alongside nocturnal activity by Geometridae moths. Camera-based observations provided higher functional resolution and revealed a different visitor assemblage than previous eDNA metabarcoding trials.

This proof-of-concept establishes a methodological foundation for forthcoming multi-day monitoring campaigns and complements ongoing studies of the mating system and population-genetic analyses.

## Introduction

Understanding how pollination dynamics vary along elevational gradients is essential for assessing potential pollinator limitation and its genetic consequences for isolated plant populations (Dai et al. 2017; Tong et al. 2021). For narrow-range endemics such as the tertiary paleoendemic cliff-dweller *Physoplexis comosa* (L.) Schur (Campanulaceae), reproductive success may be closely linked to the availability of efficient pollinators across varying abiotic conditions along a broad elevational range (200–2,800 m a.s.l.).

Although the species exhibits a specific floral morphology – characterized by permanently fused corolla lobes, basal corolla openings available at anthesis for nectar access, and secondary pollen presentation along the style – it remains unknown whether it is capable of autonomous self-pollination or relies on insect-mediated cross-pollination to set seeds.

To clarify the mating system, targeted floral isolation experiments were conducted during the 2026 flowering season, with seed set analyses planned after capsule harvest in October 2026. Establishing reliable baseline information on active floral visitor assemblages and visitation rates is therefore critical for interpreting potential pollinator limitation across the species’ elevational range (Ashman et al. 2004; Kudo and Ida 2013; Inouye 2020; Körner 2021).

In this Short Communication, we present a technical proof-of-concept for camera monitoring on vertical rock face habitats. Our long-term aim is to qualitatively and quantitatively document diurnal and nocturnal floral visitor assemblages along an elevational gradient in the Eastern Alps, providing the methodological foundation for subsequent ecological and genetic assessments.

While the robustness of floral visitor communities to climate warming represents an important ecological question, this Short Communication focuses solely on establishing the technical feasibility of camera-based monitoring in extreme terrain and evaluating its performance relative to previous eDNA metabarcoding.

## Materials&Methods

### Study Species and Study Sites

*Physoplexis comosa* (L.) Schur (Campanulaceae) is a tertiary paleoendemic cliff-dwelling species restricted to dolomitic and limestone rock faces and crevices across the Southern Limestone Alps, ranging from the Bergamasque Alps in the west to the Julian Alps in the east. To establish a methodological baseline for future investigations of floral visitor assemblages along an elevational gradient, monitoring was conducted as a pilot study to prepare a comprehensive project encompassing a broader elevational range from 200 m to 2,400 m a.s.l. Field recording spanned five representative sites from 1,480 m a.s.l. on the slopes of Monte Grappa to 2,400 m a.s.l. beneath Monte Lagazuoi (Table 1). Recording sessions were executed under stable weather conditions, characterized by clear to partly cloudy skies, light air to gentle breezes (2.2–18.7 km/h), and ambient daytime temperatures ranging between 9 °C and 21 °C. The climatic data were excerpted from the Open-Meteo API (Zippenfenig 2024).

**Table 1.** Overview of study sites, sampling dates, environmental baseline conditions, plant phenology, and applied monitoring methodologies for *Physoplexis comosa* during the 2026 field campaign.

| Location | Mte. Grappa | Mte. Altissimo | Mte. Tremalzo | Passo Falzarego | Lagazuoi Piccolo |
| --- | --- | --- | --- | --- | --- |
| Date | 026/06/2026 | 07/07/2026 | 08/07/2026 | 23-24/07/2026 | 25/07/2026 |
| Time | 8:50-10:00 | 10:15 - 11:45 | 10:00 - 11:30 | 17:00-15:00 | 09:00-11:00 |
| Duration of stay | 70 min. | 90 min | 90 min | 22 hours /overnight/ | 120 min |
| Elevation | 1400 m a.s.l. | 1930 m a.s.l. | 1880 m a.s.l. | 2100 m a.s.l. | 2400 m a.s.l. |
| Abundance /*Phys. com.*/ | Low | High | Medium-Low | Low | High |
| Temperature | 20.9-21.4 °C | 14.8-17.7 °C | 18.3-19.2 °C | 12.5-14.2 °C | 9–12.7 °C |
| Solar radiance | High | High | Medium | High | High |
| Cloud cover | Low | Low | Low | Low | Low |
| Windiness | Light air / light breeze | Light air | Low breeze | Gentle breeze | Light air |
| Bagging | NO | YES | NO | NO | NO |
| Camera recording | NO | YES | YES | YES | YES |
| Phenology | OPTIMUM | OPTIMUM | OPTIMUM | OPTIMUM / LATE | OPTIMUM |

### Camera Setup

To document diurnal and nocturnal floral visitors under steep cliff conditions, a two-tier camera monitoring protocol was applied: (i) **Diurnal Monitoring:** At each site, three Lamax 9.0 action cameras were deployed during peak diurnal insect activity (Table 1). Additionally, continuous 24-hour recording was conducted using three custom Raspberry Pi camera systems at the same sites (Fig. 1a). Cameras were mounted on tripods set up at a distance of 30–70 cm from target inflorescences (settings: 1080p resolution at 30 fps); (ii) **Night-Vision Monitoring:** at Passo Falzarego (2,100 m a.s.l.), the setup included a NoIR camera module paired with an automated 850 nm infrared LED illuminator to enable nocturnal monitoring without altering insect behaviour or inducing phototropic responses.

**Figure 1:**
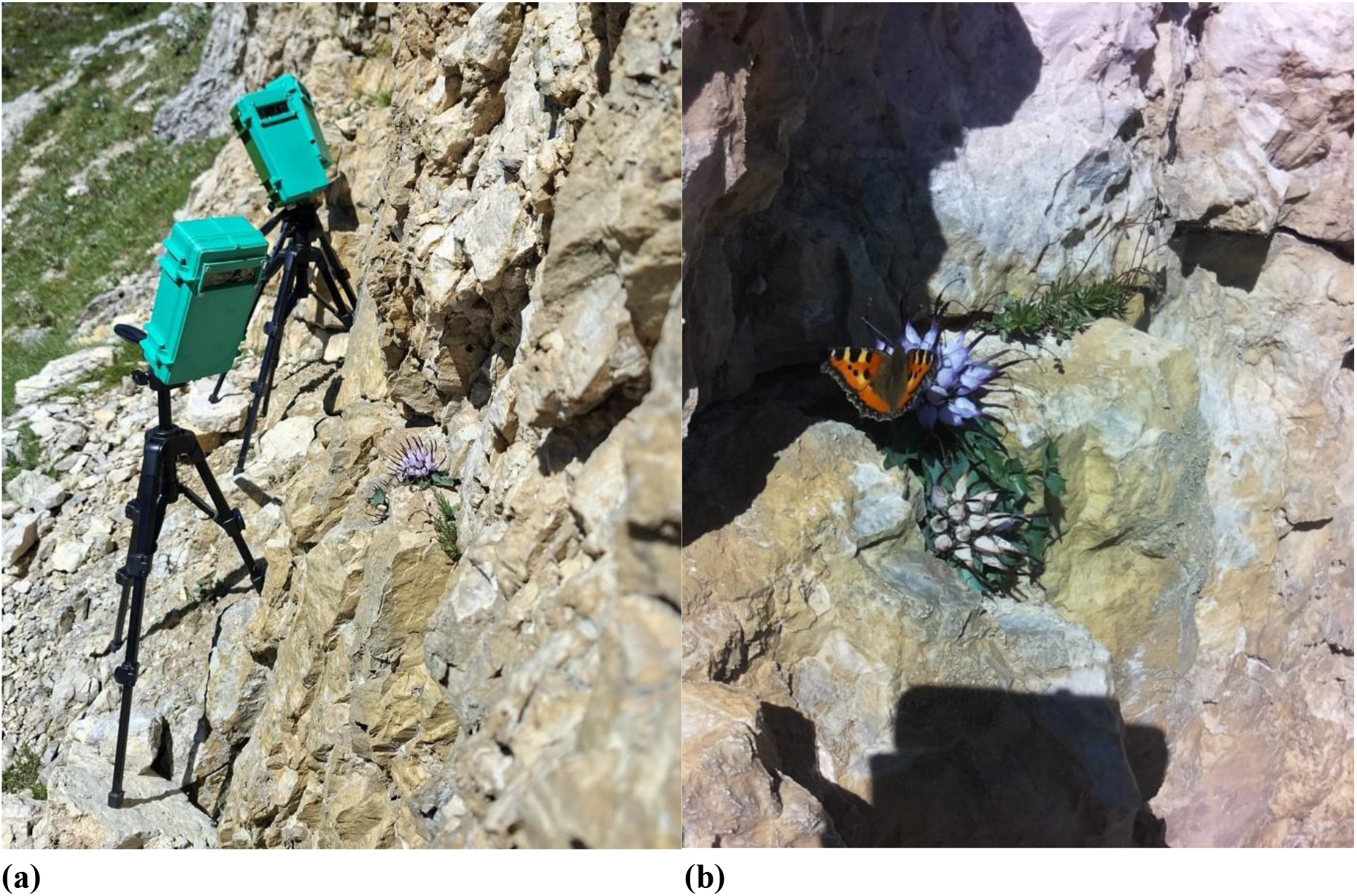
Field monitoring setup and confirmed floral visitor. **(a)** Custom Raspberry Pi-based camera setup for combined diurnal and nocturnal monitoring, deployed at Passo Falzarego (2,100 m a.s.l.). Three camera units are directed at inflorescences on a steep dolomitic rock face adjacent to blooming *Physoplexis comosa*. **(b)** Diurnal visitor contact event: *Aglais urticae* visiting an inflorescence of *P. comosa* at the same study site.

### Data Processing, Visitor Annotation, and Data Analysis

To optimize data processing from high-resolution raw footage, video files were downsampled into single image sequences using FFmpeg at sampling intervals of 1 to 3 seconds per frame (1–0.33 fps), tailored to the respective recording setup. Shorter recordings were reviewed manually, with insect-visitation frames extracted and reassembled into consolidated clips for manual review. A valid visitation event was defined as physical contact between an insect and any floral element of the inflorescence, whereas non-contact fly-bys were excluded. Taxa were identified by the authors mainly to the family level (specifically Geometridae, Syrphidae, Muscidae, and Nymphalidae) and rarely to the genus level.

To handle extensive datasets from prolonged nocturnal monitoring (>10,000 frames), image sequences were initially processed via the AddaxAI platform (v. 7.8.0, van Lunteren 2023) using MegaDetector v5a for automated target detection. All frames flagged with automated bounding boxes were subsequently visually inspected and validated. Confirmed animal detections were then manually classified to the finest possible taxonomic level (family, genus or species), following a two-step validation protocol for floral visitors similar to Stark et al. (2023).

All daytime video and image sequences were analysed manually by the authors. To complement the camera-based recording, direct *in situ* field observations were conducted during daylight hours while monitoring the sites. Consequently, visitation data are presented separately in Table 1, distinguishing between camera-recorded visits and direct field observations.

Given the pilot nature and qualitative scope of this study, data processing was limited to descriptive evaluation. Floral-visitor taxa were compiled per site and elevational step to document basic visitor presence, assemblage composition, and diurnal versus nocturnal activity patterns. Formal statistical modelling is reserved for forthcoming multi-day screening campaigns, where standardized observation windows will allow quantitative analyses.

## Results

Continuous camera-monitoring and direct field observations revealed a diverse spectrum of floral visitors across all study sites, represented by Diptera, Hymenoptera, and Lepidoptera (Table 2).

**Table 2.** Comparison of floral visitor taxa recorded on *Physoplexis comosa* via continuous camera monitoring and direct field observations across study sites during the 2026 campaign.

| Location / Site | Elevation (m a.s.l.) | Date | Camera-recorded floral visitors | Directly observed floral visitors |
| --- | --- | --- | --- | --- |
| Monte Grappa<br>45.8622439N, 11.8087256E | 1,480 | 26/06/2026 | No cameras used | Apidae (*Bombus* sp.), Nymphalidae (*Erebia* sp., *Vanessa cardui*) |
| Monte Tremalzo<br>45.8414450N, 10.7008447E | 1,880 | 08/07/2026 | Apidae (*Xylocopa* sp.), Syrphidae | Syrphidae |
| Monte Altissimo<br>45.8059053N, 10.8886053E | 1,930 | 07/07/2026 | Apidae (*Bombus* sp.) | Apidae (*Bombus* sp.) |
| Passo Falzarego<br>46.5237656N, 12.0228939E | 2,100 | 24/07/2026 | Syrphidae, Muscidae, Geometridae, Nymphalidae (*Aglais urticae*, *Erebia* sp.) | Syrphidae, Muscidae |
| Monte Lagazuoi Piccolo<br>46.5273619N, 12.0159686E | 2,400 | 25/07/2026 | Muscidae | Muscidae |

Across the five study locations (1,480–2,400 m a.s.l.), automated camera monitoring was successfully conducted at four sites, while the lowest elevation site (slopes of Monte Grappa, 1,480 m a.s.l.) was assessed solely via direct in situ observations (Table 1). Lower to subalpine sites (1,480–1,930 m a.s.l.) were characterized by active diurnal foraging of Hymenoptera/Apidae (*Bombus* sp., *Xylocopa* sp.) and Lepidoptera/Nymphalidae (*Erebia* sp., *Vanessa cardui*). At high-elevation sites (2,100–2,400 m a.s.l.), Diptera (Syrphidae, Muscidae) dominated diurnal visits alongside Lepidoptera (*Erebia* sp. and *Aglais urticae*; Fig. 1b). Automated nocturnal recording at Passo Falzarego (2,100 m a.s.l.) captured night-active Geometridae moths actively visiting the inflorescences.

The camera recordings successfully captured flower visits at four key sites (e.g., *Bombus* sp. at Monte Altissimo di Nago; Syrphidae and Muscidae at Monte Tremalzo, Passo Falzarego, and Monte Lagazuoi) while additionally capturing larger transient visitors such as *Xylocopa* sp. and various Lepidoptera (Table 2). Direct in situ observations carried out in 2026 recorded visiting insects across all five sampled locations, validating camera recordings. These records included major functional floral visitors such as *Bombus* sp. and *Vanessa cardui* at Monte Grappa, as well as various Syrphidae and Muscidae across higher-located sites (Monte Tremalzo, Passo Falzarego, Monte Lagazuoi).

These records complement preliminary direct field observations from 2023 (K. Iberl, pers. obs.), which documented sporadic visits by Formicidae at Monte Verzegnis (Carnic Prealps), Miridae near Predil (Julian Alps), and Geometridae at Monte Altissimo di Nago (Garda Prealps).

## Discussion & Outlook

Consistent with previous studies across different environments (van Klink et al. 2024; Darras et al. 2024), camera-based monitoring provides clear and reliable functional resolution in the exposed terrain of vertical rock cliffs. Our observations suggest possible elevational shifts in diurnal floral visitor assemblages, alongside a notable nocturnal component represented by Geometridae moths at the subalpine-alpine boundary elevations (Passo Falzarego, 2,100 m a.s.l.). This activity highlights a previously underestimated nocturnal interaction pathway in the reproductive biology of this paleoendemic cliff-dweller. While plant–pollinator studies have historically focused on diurnal taxa such as Apidae and Syrphidae (e.g., Walton et al. 2020), nocturnal Lepidoptera may provide functional complementarity and resilience in harsh, high-elevation environments (compare Macgregor and Scott-Brown 2020).

At the same time, specific environmental constraints should be kept in mind when using camera setups to monitor floral visitors on exposed rock faces. While continuous camera monitoring (Droissart et al. 2021) effectively captures visitation events in steep habitats, computer-vision workflows (e.g., MegaDetector embedded in Addax) depend on stable frames to prevent motion-driven detection artifacts, such as those theoretically caused by wind-driven plant motion. In our pilot deployment, the pipeline performed reliably in isolating insect visitation events, though detection sensitivity was influenced by confidence thresholds. Specifically, applying a 15% detection threshold in Addax led to a small number of missed visits, which were subsequently identified during manual cross-validation.

Beyond potential technical challenges under adverse weather, biotic flight thresholds of diurnal alpine Diptera and Hymenoptera can lead to “false-zero” visitation records when ambient temperatures drop below 10–12°C or during dense cloud cover (compare McCall and Primack 1992; Körner 2021). However, morning records of Muscidae at the highest site (slopes of Lagazuoi Piccolo, 9–12.7 °C) suggest that direct solar radiation on east-facing cliff faces can locally overcome low ambient air temperatures by warming the rock microenvironment and ectothermic visitors. Continuous camera monitoring in cliff environments is therefore methodologically most informative when interpreted alongside baseline abiotic conditions and microclimatic exposure (Table 1).

The necessity for a visual monitoring approach is further underscored by initial methodological comparisons using environmental DNA from floral surfaces (eDNA): while eDNA metabarcoding offers promising, non-invasive monitoring options, preliminary testing in our study system revealed that floral surface DNA was dominated by resident micro-arthropods (e.g., Thysanoptera, Acari), whereas transient flying visitors (Hymenoptera, Diptera) went largely undetected (Kirschner et al. in prep.). Consequently, continuous camera monitoring was required to reliably record the active pollinator assemblage of *Physoplexis comosa*. However, while this pilot campaign confirms the technical feasibility of both diurnal and nocturnal camera monitoring on vertical rock faces, capturing the full spatio-temporal variation in visitor assemblages across the species’ broad elevational gradient requires extended observation windows. A comprehensive quantitative field study along the entire elevational gradient is therefore planned for the upcoming season.

## Acknowledgements

We thank the Nature Conservation Authority of the Autonomous Province of Trento (Servizio Sviluppo Sostenibile e Aree Protette) for granting research permits (protocol no. S175/17.11.3-2026-5/58 E/VF). We gratefully acknowledge the field work support provided by student assistants Laura Hraj and Kateřina Příbramská. We are also grateful to Dr. Rudolf Ritt (Chair, Naturwissenschaftlicher Verein Passau e. V.) for his taxonomic verification of floral visitor observations in 2023.

